# Reprogramming of 3D genome structure underlying HSPC development in zebrafish

**DOI:** 10.1101/2024.01.06.574496

**Authors:** Min He, Xiaoli Li, Bingxiang Xu, Yibo Lu, Huakai Liu, Ziyang An, Wenqing Zhang, Feifei Li

**Author notes:** Correspondence (FFL) and (WQZ). These authors contributed equally to this work (Min He, Xiaoli Li).

## Abstract

Development of hematopoietic stem and progenitor cells (HSPC) is a multi-staged complex process that conserved between zebrafish and mammals; however, the mechanism underlying HSPC development is not fully understood. Chromatin conformation plays important roles in transcriptional regulation and cell fate decision, its dynamic and role in HSPC development is poorly investigated. Here, we performed chromatin structure and multi-omics dissection across different stages of HSPC developmental trajectory in zebrafish. Chromatin organization of zebrafish HSPC resemble mammalian cells with similar hierarchical structure and characteristics. We revealed the multi-scale reorganization of 3D genome and its influence on transcriptional regulation and transition of cell function during HSPC development. Nascent HSPC is featured by loose conformation with obscure structure at all layers. Notably, PU.1 was identified as a potential factor mediating formation of promoter-involved loops and regulating gene expression as well as HSPC function. Our results provided a global view of chromatin structure dynamics associated with development of zebrafish HSPC and discovered key transcription factor involved in HSPC chromatin interactions, which will provide new insights into the epigenetic regulatory mechanisms underlying vertebrate HSPC fate decision.

## Introduction

Understanding the regulatory mechanism underlying HSPC fate determination at different developmental stages is a primary goal of hematopoiesis biology. This is helpful in improving generation of functional HSPC both in vivo and in vitro. The HSPC development process is highly conserved between zebrafish and mammals and a series of important findings of HSPC ontology are based on zebrafish[1, 2]. For example, HSPC generation through endothelial-to-hematopoietic transition (EHT) is directly observed in zebrafish embryos[3]. There are three waves of hematopoiesis during zebrafish or mammalian development, with nascent HSPC arising from the ventral wall of dorsal aorta (DA) of zebrafish or aorta-gonad-mesonephros (AGM) region of mammals through the process of EHT, acquiring the ability of self-renewal and reconstruction of all blood lineages[4]. Then this group of cells move to caudal hematopoietic tissue (CHT) of zebrafish or fetal liver of mammals to be fetal HSPC which can rapid expansion and differentiation[5, 6]. Finally, these cells seed into kidney marrow (KM) of zebrafish or bone marrow of mammals, to become adult HSPC and support adult hematopoiesis[7].

Although significant achievements have been made to know this process, a comprehensive understanding of the dynamic regulatory mechanisms governing HSPC development is still lacking. Recent studies showed that despite the critical role of transcription factors (TFs), epigenetic modifications are also important in HSPC fate decision[8, 9]. Chromatin conformation is fundamental for transcriptional regulation via multiple mechanisms, from long-distance interactions between enhancers and promoters to higher-order chromosome compartments and topological associated domains (TADs) that can act as transcription restrained units[10, 11]. Recent studies have shown 3D genome rearrangement participate in hematopoietic differentiation and disease[12–14]. Role of chromatin conformation on HSPC development have preliminary explored in mice, which deepened our understanding of mechanism on HSPC fate determination [15, 16]. However, the development of HSPC is a multi-staged complex process, whether the regulatory role of 3D genome to HSPC development is conserved among vertebrate and what factors participating in regulation of HSPC development through 3D genome needs further investigation.

Here, we use zebrafish as a hematopoietic development model organism to investigate the dynamic changes in chromatin configuration during HSPC development. Multi-omics data including sisHi-C, H3K27ac ChIP-seq, ATAC-seq and RNA-seq were generated to comprehensively dissect 3D genome rearrangement and its relation to transcriptional changes of zebrafish HSPC. We found that the features of 3D genome of zebrafish HSPCs are highly similar to those observed in mammals. The development of zebrafish HSPC is accompanied by reprogramming of all layers of chromatin structure. In particular, the structural strength of nascent HSPC is much weaker. In addition, a series of transcription factors were identified potentially involved in mediating promoter-enhancer interactions and regulating HSPC development. Our study contributes to a deeper understanding of the epigenetic regulatory mechanisms underlying vertebrate HSPC development.

## Results

### Adult HSPC of zebrafish exhibits hierarchical chromatin structure similar to mammalian cells

In order to reveal the chromatin structure of zebrafish HSPCs, we first performed four replicates of sisHi-C on FACS sorted cd41+ gata1-adult HSPCs from three-month-old transgenic zebrafish Tg(cd41:GFP gata1:DsRed). RNA-seq, ATAC-seq and H3K27ac ChIP-seq were also conducted to illustrate the characteristic of zebrafish HSPC chromatin folding (Fig 1A). A total of 172842225 valid pairs were obtained from the four Hi-C replicates. GenomeDisco analysis showed replication score of any two replicates are higher than 0.85 at both 50kb and 100kb resolution, so we combined the four replicates in the following analysis (Fig S1A). The Hi-C contact map of zebrafish adult HSPC showed canonical hierarchical chromatin organization at different resolutions, including compartments, TAD and loops, similar to mammalian cells (Fig 1B).

**Figure 1.**
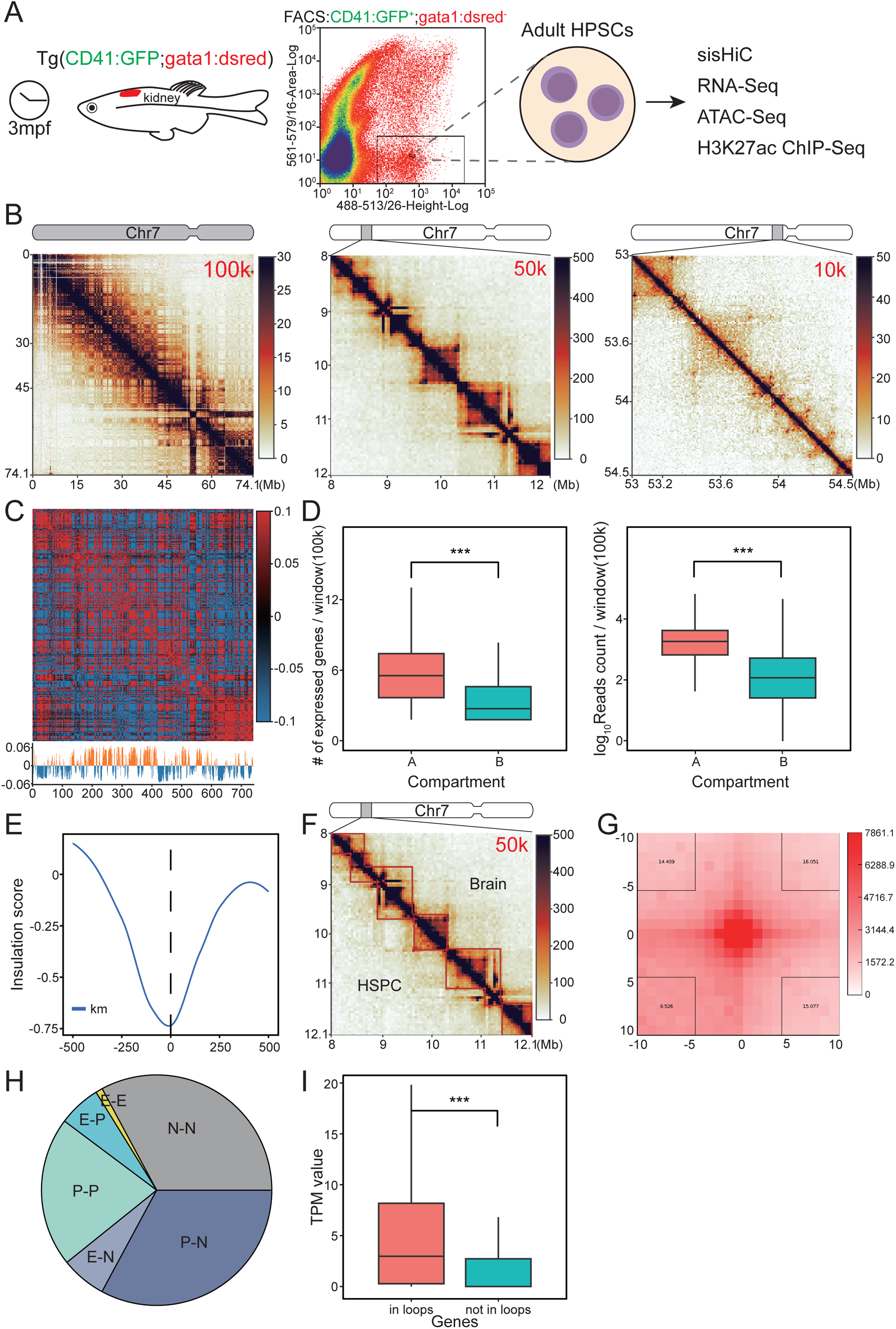
Characteristics of zebrafish HSPC 3D genome organization. (A) Schematic diagram of experimental design. (B) Hi-C contact matrix of chromosome 7 at 100kb, 50kb and 10kb resolution showed as example. (C) Chromatin compartmentalization of chromosome 7. The autocorrelation matrices and the first eigenvector profiles are shown. In the first eigenvector, compartment B is colored as blue and A as orange. (D) Boxplot showing the distribution of gene density and RNA-seq reads density in the A/B compartment. (E) Genome-wide insulation score profiles around TAD boundaries. (F) TADs detected in 8-12Mb region of chromosome 7 in both HSPC and brain are shown as an example. (G) Aggregate loop plots showing the strength of interactions between HSPC loop anchors. (H) The distribution of identified interactions. E, enhancer; P, promoter; N, neither enhancer nor promoter. (I) Distribution of transcript per million (TPM) expression value of genes involved and not involved in loops. ***, p<0.001.

At the compartment level, the autocorrelation matrix exhibits classic plaid pattern, and almost half of the genomic regions were assigned as A and B compartments, respectively (Fig 1C). Integrating with RNA-seq data, we found that A compartments contained more expressed genes (p< 2.22e-16, wilcox.test) and the expression level of encompassed genes was also higher compared with B compartments (p< 2.22e-16, wilcox.test, Fig 1D). Similarly, A compartment showed more enrichment of H3K27ac ChIP-seq peaks and ATAC-seq open regions comparing with B compartments (p< 2.22e-16 and < 2.22e-16, wilcox.test, respectively, Fig S1B). These results showed that A compartments are more active in zebrafish HSPC similar to mammalian cells. At 50kb resolution, a total of 1643 TADs with a median size of 800 kb were identified in adult HSPC. The accuracy of called TADs was verified by the strongest insulation of aggregated boundaries (Fig 1E). Similar to mammalian cells, the TAD boundaries of zebrafish HSPC enriched for transcribed TSS (Fig S1C). In order to illustrate the conservation of TAD structures between tissues, we analyzed publicly available Hi-C data of zebrafish brain (GSE134055)[17] and detected 1595 TADs. Of these, 1175 TADs are shared between brain and HSPC, accounting for over 70% of the total TADs in both tissues. This ratio almost approached the overlap ratio of biological replicates of HSPC (Fig S1D). Taking 8Mb to 12Mb region of chromosome 7 as an example, the TAD structure between adult HSPCs and the brain is highly similar (Fig. 1F). Although most majority of TADs are conserved, there are still 420 brain-and 468 adult HSPC-specific TAD boundaries. We analyze the function of genes located in tissue-specific TAD boundaries and found that enriched pathways are related to the function of specific tissues. For example, phospholipid metabolic process which is important for brain function is most enriched in brain specific boundaries (Fig S1E). While the enriched pathway in HSPC is rhythmic process which has been reported to play crucial role in hematopoietic development (Fig S1E)[18]. These results showed that TADs are highly conserved between tissues in zebrafish and the specific boundaries are correlated with tissue function. Finally, at the loop level, a total of 4189 loops were detected in adult HSPC using hiccups. The aggregated peak analysis (APA) showed enrichment of interaction, indicating high confidence of identified loops (Fig. 1G). In order to characterize the function of Hi-C loops, we identified distal enhancers genome-widely taking advantage of H3K27ac chip-seq data, and loops were assigned to functional elements. We found that most majority of identified loops connect enhancer (E) or promoter (P) (Fig. 1H). In addition, expression of genes involved in loops are significantly higher than genes not connected by loops, implying the functionality of detected loops (p< 2.22e-16, wilcox.test, Fig. 1I).

In summary, we conducted the first investigation into the chromatin conformation of zebrafish HSPC and discovered a hierarchical organization, including compartments, TADs and loops. In addition, the features of different levels of chromatin structure of zebrafish HSPCs are highly similar to those observed in mammalian cells.

### Dispersed chromatin structure in zebrafish nascent HSPC

Next, we want to reveal the reorganization of chromatin structure and its contribution to the development of zebrafish HSPC. Nascent HSPCs from the AGM region at 36 hpf, as well as fetal HSPCs from the CHT region at 3 dpf of transgenic zebrafish Tg(cd41:GFP gata1:DsRed) were collected and performed sisHi-C for at least two replicates (Fig 2A). A total of 26350389 and 118555641 valid pairs were obtained for nascent and fetal HSPC, respectively (Table S1). High reproducibility of Hi-C experiments was validated by a median GenomeDISCO score near 0.8 for all replicates (Fig S2A). The valid pairs and replication score of nascent HSPC was relatively attenuated comparted with fetal HSPC, which may due to the rarity and relatively low number of nascent cells. To make the Hi-C data of different stages comparable, we downsampled the pooled valid pairs to the number of nascent HSPC which had the lowest valid contacts. Generally, Hi-C contact maps are more similar between fetal HSPC and adult HSPC, but substantially different from nascent HSPC (Fig 2B). Median GenomoDisco score of Hi-C contact maps were 0.844 and 0.904 comparing fetal HSPC with nascent and adult HSPC, respectively (Fig S2B). Visually, interactions in nascent HSPC are concentrated near diagonal area, while more long-range interactions spanning dozens of megabases were observed in fetal and adult HSPC. This trend was obvious when subtracting contact matrix of fetal and adult HSPC by the matrix of nascent stage (Fig 2C). Contact frequency decay curves also showed this trend, with contacts ranging from approximately 1 Mb to 25 Mb were much lower in nascent HSPC compared with fetal and adult HSPC (Fig 2D). Using Jensen–Shannon divergence (JSD) to measure the closeness of two decay curves, JSD was 0.0017 and 0.0012 comparing fetal HSPC with nascent and adult HSPC, respectively. In order to further clarify these, we calculated frequency of intra-chromosomal contacts with different distance, that is smaller than 1Mb, 1-25Mb and larger than 25Mb, in the three stages of HSPC. Result also showed that nascent HPSC are depleted with interactions with length of 1-25Mb, which may impair megabase-sized modular structure, such as TADs (Fig S2C).

**Figure 2.**
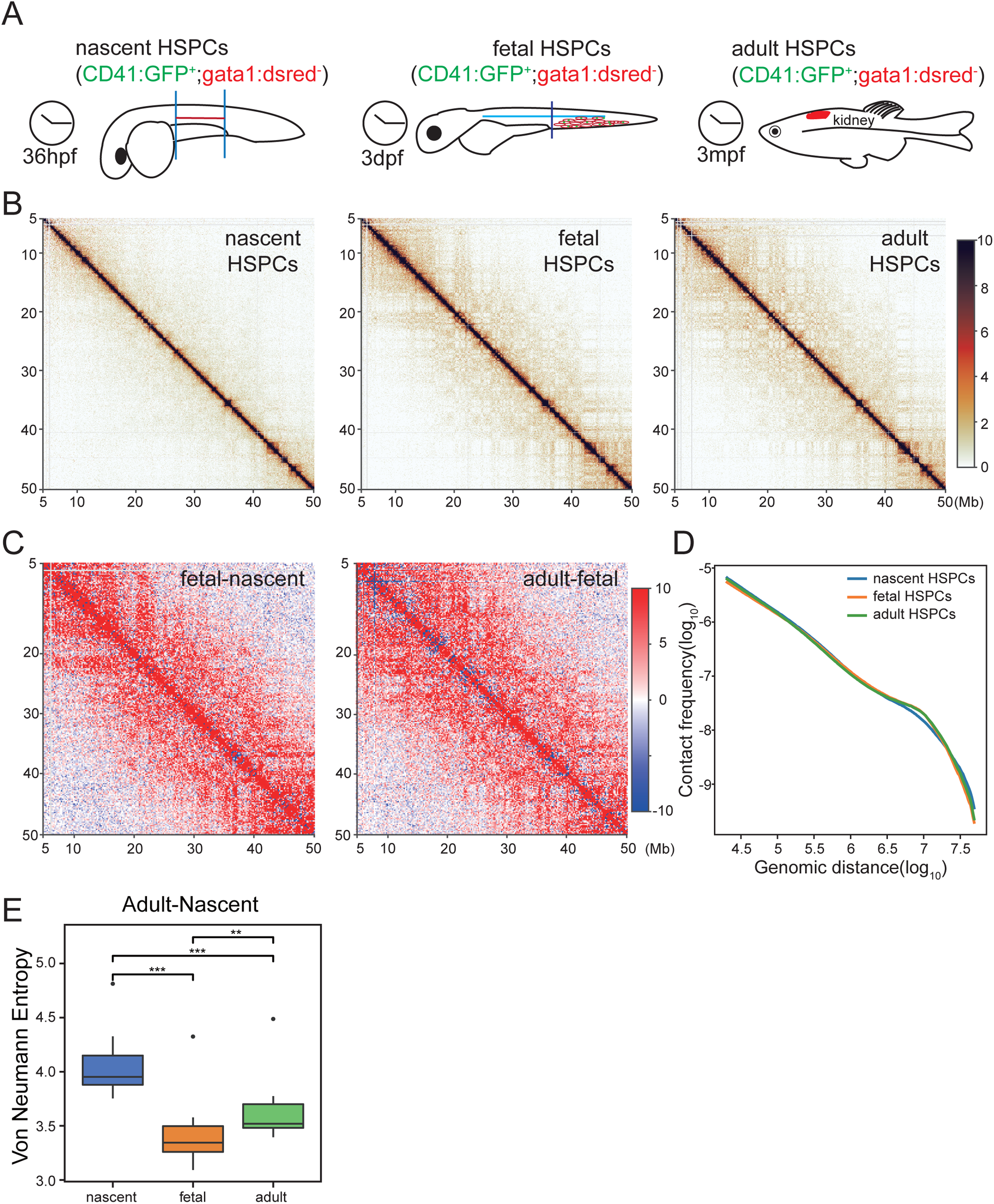
Global reorganization of chromatin structure during zebrafish HSPC development. (A) Schematic representation of chromatin conformation detection for nascent HSPCs in the AGM region at 36hpf and fetal HSPCs in the CHT region at 3dpf. (B) A 45Mb region of chromosome 7 is shown with 50-kb resolution as an example of the contact maps during the process of HSPC development. (C) Contact heatmaps of fetal and adult HSPC were subtracted by that of nascent HSPC for the same region as B. (D) Contact frequency decay curves at different stages of HSPC development. (E) Quantification of the intra-chromosomal disorder in chromatin structure using Von Neumann Entropy (VNE). One dot represents one chromosome.

The depletion of megabase-size contacts and increased super long-range contacts in nascent HSPC reminded a more relaxed chromatin organization[12]. In addition, higher proportion of inter-chromosome interaction of nascent HSPC also indicated loose structure (Table S1). In order to investigate the folding characteristic, we calculated chromosomal level Von Neumann Entropy (VNE) index to quantify chromatin disorder (Fig 2E). The result showed that the entropy of nascent HSPC was significantly higher than that of fetal and adult, indicating more disordered chromatin structures. Then, we want to ask whether transcriptional and other epigenetic data support the more relaxed structure of nascent HSPC. RNA-seq and ATAC-seq data of adult HSPC were compared with publicly downloaded nascent and fetal HSPC data (CRA001858)[19]. Principal component analysis showed that gene expression is clearly distinguished among different stages of HSPC development (Fig S2D). A total of 6712 and 6544 differential expressed genes were identified between two consecutive stages (Fig S2E). Clustering analysis showed that expression pattern of adult HSPC and fetal HSPC are more similar than nascent HSPC (Fig S2F). Importantly, the number of significantly downregulated genes is more than twice of upregulated genes from nascent to fetal HSPC. Similarly, chromatin openness was also different among HSPC developmental stages, and the difference was more pronounced between nascent and fetal HSPC (Fig S2G). Quantitatively, 10092 and 5693 differential open regions were identified for fetal HSPC compared with nascent and adult HSPC, respectively (Fig S2H). Similarly, nearly two folds of regions become closed than that become accessible from nascent to fetal HSPC. These results showed that transcription and chromatin availability were consistent with chromatin structure and support relaxed conformation of nascent HSPC.

In conclusion, gene transcription, chromatin accessibility and 3D genome structure were dynamic changed during zebrafish HSPC development, especially from nascent to fetal stages. Chromatin of nascent HSPC was more relaxed and relatively depleted with interactions at megabase size.

### Rearrangement of compartments affect gene expression during zebrafish HSPC development

We compared the compartment status of the three HSPC stages at 100kb resolution. Visual inspection showed that the Pearson autocorrelation matrix for nascent HSPC appears relatively obscure, and the alternating pattern of A/B compartments is less pronounced compared with fetal and adult HSPC (Fig 3A). We first investigate the switching of A/B compartments positions during HSPC development. About 18% and 12% genomic regions undergo compartment changes from nascent to fetal HSPC and from fetal to adult HSPC, respectively (Fig S3A). Genes in A compartment had higher transcriptional level than B compartments for all stages (Fig S3B). In addition, expression of genes contained within regions changing from A to B compartment were decreased, while gene expression in B to A switched regions are increased, which further verified the active characteristics of A compartment (Fig 3B). Genes located in the switch regions are correlated with characteristic of HSPC stages. For example, Genes switched from B to A compartment and showed transcriptional upregulation from nascent to fetal HSPC were enriched in pathways such as ‘RNA processing’ and ‘ribosome biogenesis’ (Fig 3C), while from fetal to adult HSPC, ‘lipid biosynthetic’ and ‘phagocytosis’ related pathways are enriched (Fig S3C), in accordance with the rapid proliferation of fetal HSPC and adaptive immunity of adult HSPC. We detected some important regulatory factors switched compartments during HSPC development. For example, runx3 was previously reported to contribute to the maintenance of HSCs in fetal liver, but play no role at the onset of definitive hematopoiesis[20]. It located in B compartment in nascent HSPC with low gene expression and chromatin accessibility, while changed to A compartment with higher expression and accessibility in fetal HSPC (Fig 3D). These results showed that assignment of compartments changed during HSPC development, especially from nascent to fetal HSPC. In addition, the switching of A/B compartments is associated with changes of gene expression and function of specific stages of HSPC.

**Figure 3.**
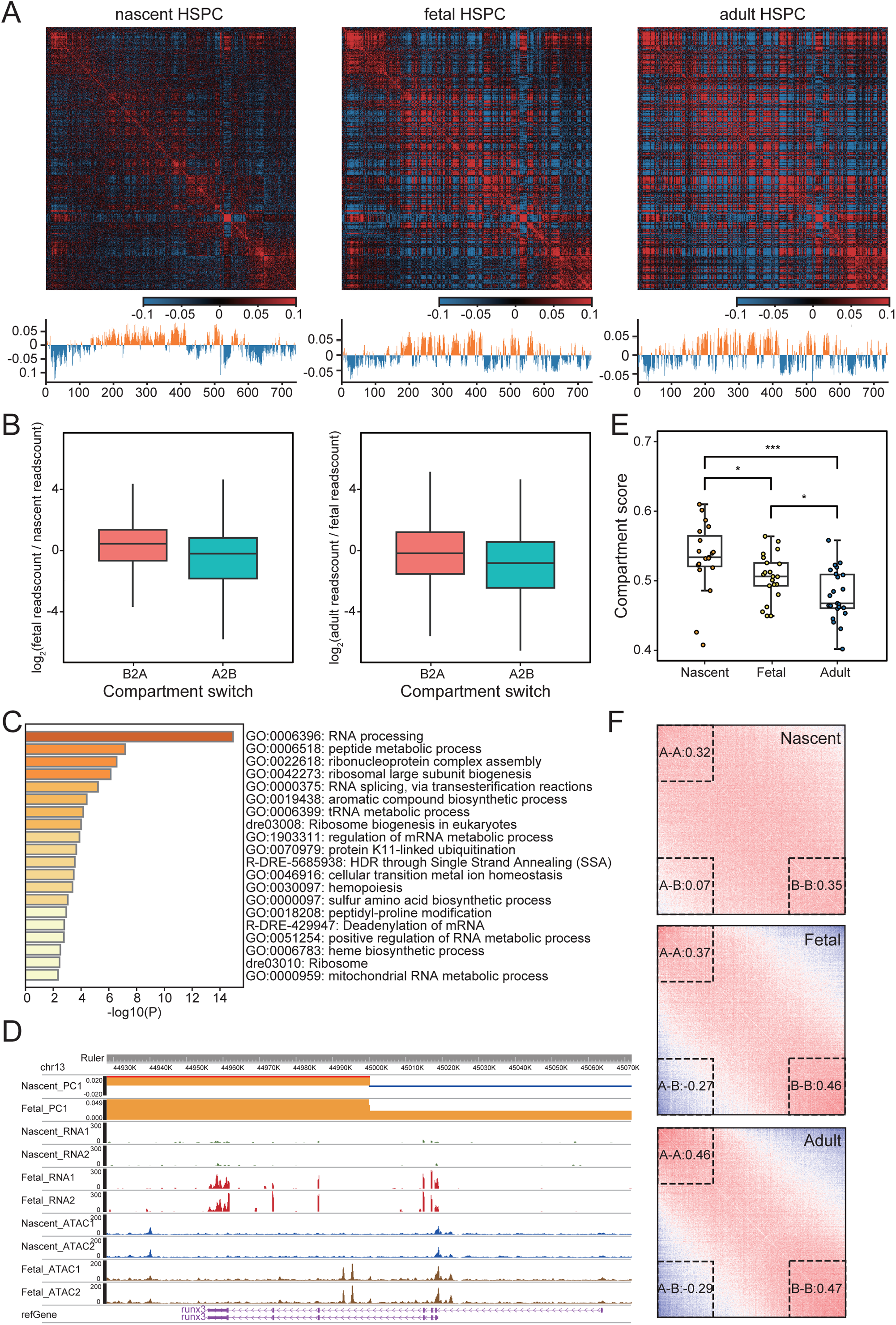
Compartmentalization reprogramming associated with expression changes during HSPC development. (A) Chromatin compartmentalization at the three developmental stages. Chromosome 7 is shown as an example as in Figure 1C but with down-sampled data. The autocorrelation matrices and the first eigenvector profiles are shown. (B) Expression changes of genes located within compartment switching regions. (C) Enriched pathways of genes located in B2A switch region and upregulated from nascent to fetal HSPC. (D) Switching of compartment assignment near runx3 gene is shown by first eigenvector profiles. ATAC-Seq and RNA-Seq signal for both replicates are displayed in the rest of the tracks. (E) The chromosome-wise compartment scores at each developmental stage. (F) Compartmentalization saddle plots of different stages during HSPC development. Numbers represent relative interaction strength between compartments.

To further determine whether changes in compartmentalization occur coincidently with local changes in chromatin accessibility, we compared compartment assignment with ATAC-seq data. A compartment had higher chromatin accessibility than B compartments in all stages (Fig S3D) and the stage-specific ATAC peaks mainly occurred in stage-restricted compartment A regions (Fig S3E and S3F). These indicate the coordination of large-scale compartment and local chromatin accessibility. In addition, we found that nascent and fetal specific ATAC-seq peaks were most enriched in motifs corresponding to c-JUN and ETV2 (Fig S3G), which are known to be involved in HSPC emergence and proliferation, respectively[21, 22]. However, the most enriched transcription factor is CTCF in adult HSPC specific peaks, which implying the formation of more loops in adult HSPC.

Next, we compared the strength of compartmentalization at the three HSPC stages. Compartment scores (AB / AA + BB) were calculated and the result showed that score of nascent HSPC is significantly higher than fetal HSPC, indicating weaker compartmentalization in nascent HSPC (p = 0.015, wilcox.test, Fig 3E). Compartment score of fetal HSPC was also significantly higher than adult HSPC, implying the gradually increasing separation of A and B compartments during HSPC development (p = 0.033, wilcox.test, Fig 3E). Saddle plot showed the same trend with gradually higher segregation levels during HSPC development, especially form nascent to fetal stage (Fig 3F). The above results indicate that rearrangements of compartmental structures is associated with changes of gene expression and HSPC function during zebrafish HSPC developmental process.

### TAD structure kept stable but much weaker in zebrafish nascent HSPC

We then investigate changes of chromatin TAD structure during zebrafish HSPC development. At a resolution of 50Kb, a total of 1760 and 1667 TADs were identified with median length of about 750 kb and 800 kb in nascent and fetal HSPC, respectively. Accuracy of TADs were verified by the lowest IS at aggregated TAD boundaries. The position of TADs remained largely unshifted during HSPC development. There are 1044 TADs were shared by all three stages, which is similar to that observed between adult HSPC and brain (Fig. 4A, Fig S1E). In addition, when comparing two consecutive stages, about 60% TAD boundaries remained stable and not shifting more than one bin. This proportion is similar to that observed between two biological replicates (Fig S4A). Fig S4B showed a region containing four TADs that kept absolutely stable during HSPC development. These results showed that TAD position largely kept stable during HSPC development.

**Figure 4.**
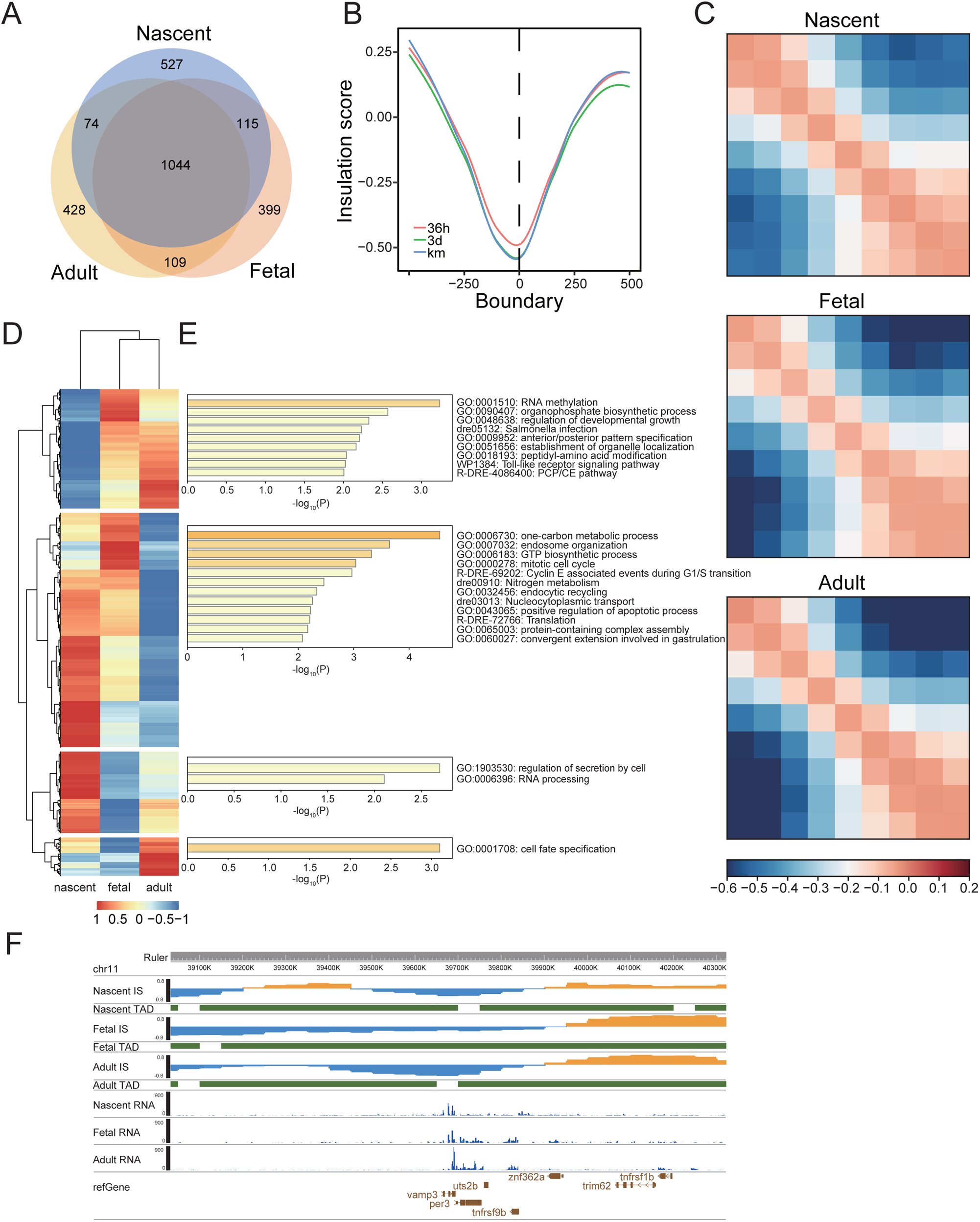
TAD structure kept stable but much weaker in zebrafish nascent HSPC. (A) Venn graphs showing the overlap of TADs between different stages of HSPC. (B) Genome-wide insulation score profiles around TAD boundaries. (C) Observed/expected (O/E) aggregate plot of interaction profile centered on TAD boundaries for the three stages of HSPC. (D) Heatmap showing the clustering of top 1000 variable TAD boundaries based on insulation score. Each row was one TAD boundary. (E) Gene ontology analysis for genes located within TAD boundary clusters. (F) Insulation score, TAD structure as well as RNA-seq signal was shown near vamp3 gene for nascent and fetal HSPC.

Then, we compared TAD strength of the three stages of HSPC and found strength are much weaker in nascent HSPC. Insulation score of TAD boundaries in fetal HSPC is comparable with adult HSPC, with mean IS of −0.530 and −0.527, respectively (p=0.92, paired t-test, Fig 4B). However, IS of nascent HSPC is much higher than fetal HSPC (mean IS = −0.478, p=0.07, paired t-test, Fig 4B). In addition, compared with nascent HSPC, fetal and adult HSPC showed more increased intra-TAD interactions and more decreased inter-TAD interaction in the stacked interaction profile centered on TAD boundaries (Fig 4C). In order to dissect the influence of these variable boundaries, we identified the top 1000 highly variable boundaries based on IS among the nearly 4000 bins that were used as TAD boundaries in any stage. These boundaries can be clustered into four categories, which corresponding to the three developmental stages specific boundaries and common boundaries of nascent and fetal stages (Fig 4D). Gene ontology analysis showed that genes encompassed in these stage-biased boundaries are associated with function of specific stage (Fig 4E). For example, cluster 2 showed active endocytic related pathways, consistent with adult HSPC function of adaptive immunity. Although overall expression level of genes in stage-specific boundaries have no significant change during development (Fig S4C), we observed some differential expressed genes associated with boundary changes. For example, up-regulation of genes VAMP3, which is associated with endocytosis and facilitates membrane fusion [23, 24], is accompanied by decreased insulation score and new TAD boundary establishment in adult HSPC (Fig 4F).

Taken together, TADs largely kept stable during HSPC development but much weaker at nascent stage in zebrafish.

### PU.1 mediate promoter-involved looping interactions in adult HSPC

We compared the loop structure of fetal and adult HSPC, due to the relatively limited valid pairs impaired loop detection of nascent HSPCs. A total of 1395 loops were identified in fetal HSPC at the resolution of 10kb. Accuracy of the called loops were validated by the high interaction frequency at aggregated peak analysis (APA) analysis (Fig 5A). A total of 1218 loops were shared (with both anchors shifting no more than one bin) between fetal and adult HSPC, implying the most majority of fetal loops are preserved at adult stage (Fig 5B). Further quantitative analysis showed the alteration of loops is actually minor. We plot the APA profile of the 2971 adult HSPC-specific loops using nascent and fetal contact matrix, and found that interactions between anchors of these loops are also higher than neighboring regions in both nascent and fetal stages (Fig S5A). In addition, cosine similarity of contact frequencies of these 2971 loops were as high as 0.87 and 0.93 when comparing nascent with fetal HSPC and fetal with adult HSPC, respectively. These observations indicated that adult HSPC specific contacts actually had been concentrated in nascent and fetal stages, which is further strengthened at adult HSPC.

**Figure 5.**
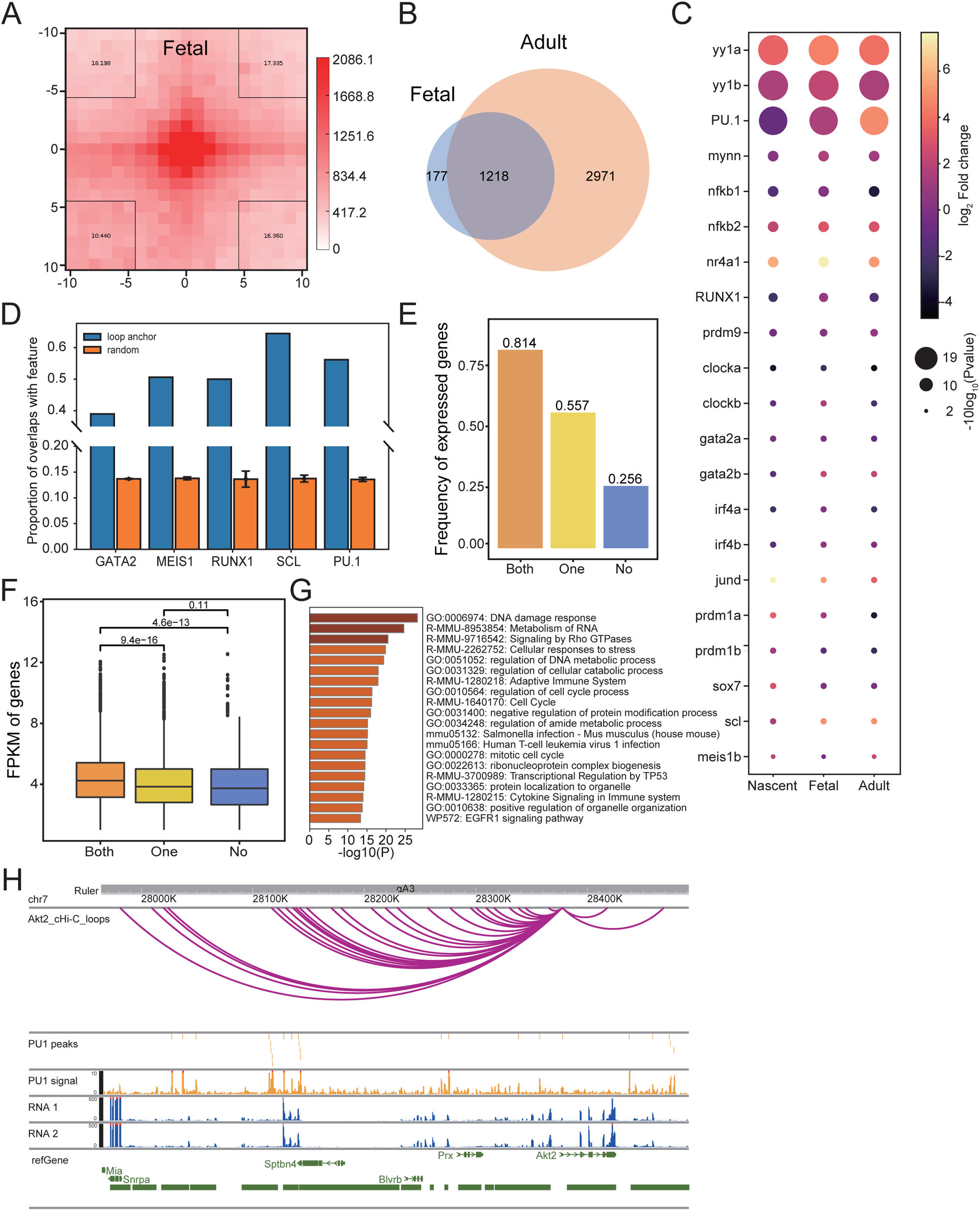
PU.1 mediate promoter involved chromatin looping in adult HSPC. (A) Aggregate loop plots showing the strength of interactions between fetal HSPC loop anchors. (B) Overlap of loops between fetal and adult HSPC. (C) Bubble plots showing gene expression and TF motif enrichment identified at H3K27ac peak region in adult HSPC loop anchors. Enriched p-value was calculated by HOMER. (D) Bar chart showing the overlap proportion of the TF-binding peaks with the loop anchors (blue) or background random selected regions (orange) for HPC7 cell. (E) Proportion of expressed genes for different groups of genes based on PU.1 occupancy on loop anchors. (F) Transcriptional level for expressed genes in different groups as in E. (G) Gene ontology analysis for genes having loops connecting gene promoter with both anchors occupied by PU.1. (H) Promoter capture Hi-C loops, PU.1 peaks and signal, as well as RNA-seq signal was present near Akt2 gene for HPC7 cell line.

Next, we focused on adult HSPC to identify transcription factor participating in mediating loop interactions. Focusing on transcriptional regulatory loops, we did motif analysis on H3K27ac ChIP-seq peak regions located in loop anchors of adult HSPC. Result showed that in addition to ETS-domain transcription factor family, such as PU.1, other 31 TFs were identified whose motifs are enriched (p < 0.01). Above results showed loop interactions of adult HSPC were already exist in nascent and fetal stages but strengthened in adult HSPC, so we paid attention to TFs that are expressed in all three stages but significantly increased in adult HSPC. The homologs of 15 TFs in zebrafish are expressed in all three stages based on RNA-seq data (Fig 5C). Several TFs with known function in HSPC development were identified, such as TAL1, RUNX1, GATA1 and PU.1[25, 26]. The most enriched TFs are YY1 and PU1. In addition, expression of PU.1 was gradually increased while YY1 transcriptional level kept relatively stable during HSPC development (Fig 5C). The result indicated that PU.1 may mediate loop interactions in zebrafish adult HSPC.

In order to further clarify the potential of identified TFs mediating loop structures and regulate transcription, we utilized a common blood stem/progenitor cell model HPC-7, which has ChIP-seq data of 5 out of the 15 candidate TFs as well as promoter-capture Hi-C (pcHi-C) and RNA-seq data[25, 27, 28] (GSE48086, GSE22178, E-MTAB-3954). We calculated the frequency of TF peaks that overlap with the interacting fragments identified by pcHi-C, and compared it with randomly picked noninteracting control regions. All of the 5 TFs showed significant enrichment at interacting regions indicating their role in genomic looping (Fig 5D). We also calculated number of loops that have specific TF binding, and found PU.1 was most frequently present on loop anchor (Fig S5B). Nearly half of promoter-involved loops have PU.1 binding on both or one anchor. To further clarify function of PU.1 in mediating chromatin loops, we analyzed influence of PU.1 binding to transcription. Genes were classified into three groups based on PU.1 occupancy on both anchors, one anchor or no anchor of loops connecting promoters. We found more proportion of genes are expressed and the gene expression level is also significantly higher in genes with both anchors binding by PU.1 than genes with one anchor binding by PU.1 (Fig 5E and Fig 5F). This difference is also obvious when comparing genes with one anchor binding by PU.1 and no anchor binding. Gene ontology analysis showed that genes with both anchors occupied by PU.1 are specially enriched in immune-related pathways, implying the role of PU.1 mediating loops in maintaining the characteristics and functions of HSPC (Fig 5G, S5C and S5D). We identified several well-known genes important for immune system having PU.1 binding on both anchors, such as Akt1 and Akt2[29] (Fig 5H and S5E).

Taken together, our results showed that PU.1 may mediate 3D genome looping interactions and regulate gene expression and function of HSPC.

## Discussion

In summary, by integrating multi-omics datasets generated from 3D genome structure, transcriptome, chromatin accessibility as well as histone modification, this study reports the structural dynamic of multi-layered 3D genome and its contribution to shape HSPC ontogeny in zebrafish which is a model animal for hematopoietic development. In particular, PU.1 was detected and verified by public data that potentially mediating chromatin loop formation and regulate gene expression as well as HSPC characteristics.

We found that chromatin of zebrafish HSPC are organized into hierarchic structure with similar feature as mammalian cells. During development of HSPC, the obscure 3D genome structure in nascent HPSC was strengthened in fetal and adult HSPC at all layers, including compartments, TADs and loops. Compared with studies in mouse, the reprogramming of chromatin structure during HSPC development have commonalities and species specificity. Murine fetal and adult HSPCs preserved large-scale compartments and TADs structure, while intra-TAD interactions are more dynamic[15]. This is in accordance with our results, which showed highly similar contact frequency decay curves as well as conserved position and strength for both compartments and TADs between zebrafish fetal and adult HSPC. The loop structure was more variable with less loops and weaker strength in fetal HSPC compared with adult HSPC. However, although murine nascent HSPC did not show impaired chromatin structure[16], our results in zebrafish nascent HSPC illustrated more relaxed chromatin organization and compromised A/B compartmentalization and TADs strength. This more disordered structure in zebrafish nascent HSPC was supported by changes in chromatin accessibility, gene expression as well as the chromatin entropy. This may underlie the substantial molecular and phenotypical differences between nascent and fetal HSPCs showed by previous studies[5, 30]. In addition, in agreement with 3D structure in early mammalian embryos is obscure but gradually enhanced during development[31], the relatively relaxed structure highlights a highly plastic state at the early stages of HSPC development and may be important for transitions from endothelial to hematopoietic properties.

The ETS-family transcription factor PU.1 is a key regulator of hematopoiesis. PU.1 is activated in HSPC and is expressed in mast cells, B cells, granulocytes, and macrophages but is switched off in T cells. Previous studies illustrate that PU.1 play crucial rules in the development of both myeloid and lymphoid lineages as well as lymphoid-primed multipotent progenitors[32–34]. For HSPC, PU.1 is important for maintenance or expansion of HSPC number in murine fetal liver[35], and for homing and long-term engraftment in the bone marrow[36]. In addition, bone marrow HSCs disrupted with PU.1 in situ could not maintain hematopoiesis and were outcompeted by normal HSCs. PU.1 also limits hematopoietic stem cell expansion and prevent exhaustion of adult HSPC[37]. These results illustrate multiple functions of PU.1 in HSPC development, maintenance and differentiation[38]. Our results provided evidence that PU.1 can regulate HSPC gene expression and function through mediating chromatin loops. Some studies have illustrated PU.1 can function as loop mediator at specific loci or genes[39, 40]. As far as we know, for the first time, our results proposed the genome-widely structural function of PU.1 in enhancer-promoter interactions in HSPC. In addition, the evidence from murine HPC7 support the conservation of PU.1 structural roles between species. One recent study in murine HSPC highlighted RUNX1 engaged in chromatin interactions and promoted hematopoiesis. Interestingly, RUNX1 and PU.1 was shown to have physical interactions[41], and the relationship of these two proteins as well as other interaction partners in mediating chromatin interactions needs further investigation. In addition, whether other identified TFs, such as YY1[42], also play roles in mediating chromatin loops in HSPC need more research.

## Materials and Methods

### Collection of Zebrafish embryos

Zebrafish strains including Tubingen, Tg(CD41:GFP), Tg(gata1:dsRed), and Tg(CD41:GFP,gata1:dsRed) were raised under standard conditions (28.5°C in system water). Zebrafish were raised, bred and staged according to the standard protocols. All experiments involving zebrafish were carried out in accordance with the guidelines set by the Institutional Animal Care and Use Committee of South China University of Technology, Guangzhou, China.

### Flow cytometry

The trunk or tail region of zebrafish 36hpf or 3dpf embryos Tg(CD41:GFP, Gata1:DsRed) was cut for collection of nascent and fetal HSPC, respectively. Before the collection of trunks, embryos of 36 hpf were dechorionated using pronase. The trunk or tail region were ground, washed by grinding fluid and filtered using 100μm cell-strainer. Then the dissociated cells were digested with dispase at 37 °C for 30 min into single cell suspension, followed by filtration using 40μm cell-strainer. The kidney marrow of 3 mpf zebrafish were filtered to be single cell suspension using 40μm cell-strainer after puffing well. Single cell suspension of different staged HSPCs were sorted and analyzed by flow cytometers MoFlo XDP (Beckman Coulter) in the purify model. The FACS data were analyzed with FlowJo software (v10, Tree star). Fluorescence markers for HSPC at different periods and regions were CD41+gata1−.

### In situ sisHi-C library preparation

The generation of in situ Hi-C library was performed as reported[43]. Briefly, cells sorted by FACS were fixed with freshly made 1% formaldehyde solution at room temperature for 10 min. 1.25M glycine solution was added to a final concentration of 0.2 M for quenching the reaction. Cells were lysed in ice-cold Hi-C lysis buffer (10 mM Tris-HCl pH 8.0, 10 mM NaCl, 0.2% Igepal CA630, 1x protease inhibitor cocktail) for 15 min. Pelleted nuclei were washed once with 1x NEBuffer 2 and incubated in 0.5% sodium dodecyl sulfate (SDS) at 62°C for 5 min. After incubating, water and Triton X-100 were added to quench the SDS. MboI restriction enzyme (NEB, R0147M) was added and chromatin was digested at 37°C for 5 hr. Biotin-14-dATP was used to mark the DNA ends followed by proximity ligation in intact nuclei. After crosslink reversal, DNA was sheared to a length of ∼300 bp with Covaris M220, then treated with the End Repair/dA-Tailing Module (NEB, E7442L) and Ligation Module (NEB, E7445L) following the operation manual. Biotin-labeled fragments were pulled down using Dynabeads MyOne Streptavidin C1 beads (Life technologies, 65001). The Hi-C libraries were amplified for 11–15 cycles with Q5 master mix (NEB, M0492L) following the operation manual. DNA was then purified with size selection. Libraries were then quantified and sequenced using NovaSeq platform (Illumina).

### ChIP-seq library preparation

ChIP-seq was conducted according to with few modifications[44]. The cells were cross-linked with a final concentration of 1% formaldehyde followed by quenching with glycine. Cells were lysed with lysis buffer (0.2% SDS;10 mM Tris -HCl, pH 8.0; 10 mM EDTA, pH 8.0; proteinase inhibitor cocktail) and sonicated to fragments about 300 - 500 bp (Bioruptor, Diagenode). Dynabeads Protein A was washed twice with ChIP Buffer (10mM Tris-HCl pH7.5, 140mM NaCl, 1mM EDTA, 0.5mM EGTA, 1% Triton X-100, 0.1% SDS, 0.1% Na-deoxycholate, Cocktail proteinase inhibitor) and was incubated with antibody at 4℃ for 2-3hours. The fragmented chromatin was transferred to the bead-antibody complex tubes and rotated at 4 °C overnight. The beads were washed once with low salt buffer (10mM Tris-HCl pH7.5, 250mM NaCl, 1mM EDTA, 0.5mM EGTA, 1% Triton X-100, 0.1% SDS, 0.1% Na-deoxycholate, Cocktail proteinase inhibitor) and twice with high salt buffer (10mM Tris-HCl pH7.5, 500mM NaCl, 1mM EDTA, 0.5mM EGTA, 1% Triton X-100, 0.1% SDS, 0.1% Na-deoxycholate, Cocktail proteinase inhibitor). After crosslink reversal, library was constructed as in-situ Hi-C.

### ATAC-seq library preparation

ATAC-seq was prepared as previously described with few modifications[45]. Briefly, 50000 fresh cells were resuspended in 50 μl of ATAC-seq resuspension buffer (RSB; 10 mM Tris-HCl pH 7.4, 10 mM NaCl, and 3 mM MgCl2) containing 0.1% NP40, 0.1% Tween-20, and 0.01% digitonin and incubated on ice for 3 min. After lysis, 1 ml of ATAC-seq RSB containing 0.1% Tween-20 (without NP40 or digitonin) was used to wash nuclei. Nuclei were resuspended in 50μl of transposition mix (10μl 5XTTBL (Vazyme TD501), 5μl TTE Mix V50, and 35μl water) and pipetted up and down 20 times to mix. Transposition reactions were incubated at 37 °C for 30 min in a thermomixer. After the tagmentation, purify sample using the Ampure XP beads. The ATAC-seq library was amplified for 11 cycles of PCR with TAE mix (Vazyme TD501) following the manual. DNA was then purified with size selection, quantified and sequenced using an Illumina sequencing platform.

### RNA-seq library construction

The mRNA library was prepared using an optimized Smart-seq2 protocol[46]. Briefly, ∼10 ng of total RNA for each sample was utilized for first-strand cDNA reverse transcription in a 30 μl RT buffer containing SuperScript II RTase (100 U), RNase inhibitor (10 U), dNTP mix (10 mM each), SS III first-strand buffer (1×), DTT (5 mM), betaine (1 M), MgCl2 (6 mM) and TSO (1 μM). The RT reaction was performed at 90min at 42 °C followed by 11 cycles of 2min at 50 °C and 2min at 42 °C for amplification and 15min at 70 °C. The full-length cDNA was subsequently amplified through semi-suppressive PCR for 17 cycles in a buffer containing 30 μl of first-strand cDNA, 37.5 μl KAPA HiFi HotStart ReadyMix (1×), ISPCR primers (0.1 μM), and nuclease-free water. The amplified full-length DNA library was purified to get rid of <500 bp fragments and other contaminants using AMPure XP beads (Beckman Coulter, A63881) following the manufacturer’s protocol. 1 ng of purified DNA was exploited for library preparation using TruePrep DNA Library Prep Kit V2 for Illumina (Vazyme). DNA was then purified with size selection. Libraries were then quantified and sequenced using Illumina platform.

### Sequencing reads pre-processing and quality control

The quality of all libraries were evaluated by FastQC (http://www.bioinformatics.babraham. ac.uk/projects/fastqc/, v.0.11.9). Reads with mean quality score less than or equal to 30 are removed. Raw reads were trimmed and removed for adapter sequences by fatsp (v.0.23.2) with paired-end default parameters. Extremely short fragment with length less than or equal to 30bp were also removed[47].

### ChIP-seq and ATAC-seq data processing

All ChIP-Seq and ATAC-seq reads were mapped to GRCz10 zebrafish genome by Bowtie2 (v.2.5.0) with very-sensitive configuration[48]. The uniquely aligned fragments with MAPQ ≥ 30 were extracted using SAMtools (v.1.9)[49]. Duplicates were removed by Picard tools MarkDuplicates (https://broadinstitute.github.io/picard/, v.2.27.4). ChIP-seq peaks were called using MACS2 with parameters “-f BMAPE -p 0.005”[50]. ATAC-seq peaks were also called using MACS2 callpeak command (v.2.2.5) with parameters “--nomodel --shift -100 --extsize 200 -q 0.05”. Peaks were first called in individual replicates. Then reads from different replicates were merged, and peak calling was performed with merged reads. Repeated peaks were then taken as those called from the merged reads that overlapped with those called in all replicates. Signal tracks were built using the bdgcmp sub-command of MACS2 with the fold enrichment over control (FE) mode. Enhancers were defined as H3K27ac ChIP-seq peaks that not overlap with 6kb region centered on transcriptional start sites. To obtain differentially accessible regions, we merged peaks from all samples to obtain a non-redundant peak set. Read pair numbers for each non-redundant peak were calculated using HTseq (v0.8.0) and compared with DESeq2 [51, 52].

### RNA-seq data analysis

After filtering, high-quality reads were aligned to GRCz10 zebrafish genome using HISAT2 with default parameters[53]. FeatureCounts was used to quantify gene expression and obtain reads count. Fold changes in gene transcription levels were estimated using DESeq2[54]. Enrichment analysis of gene function was performed in the Metascape platform (http://metascape.prg/gp/index.html).

### Hi-C data processing and visualization

HiC-Pro (v.3.1.0) was used for the processing of Hi-C data[55]. Only uniquely mapped read pairs with mapping quality no less than 10 were saved for further analysis, and dangling end reads, self-circled reads, religated reads were all trimmed out. Non-duplicated reads were used to generate Hi-C contact matrices at the binning resolution of 10kb, 50kb and 100 kb. To validate the reproducibility of data, we calculated the GenomeDisco score between two libraries[56]. Contact heatmaps were generated with matrices at different resolutions by fanc (v.0.9.25)[57]. The p(s)-curves were calculated from genome distances of 20 kb to 50 Mb separated into 500 bins logarithmically. We applied the Von Neumann Entropy (VNE) approach to quantify the disorder of chromatin structure for 100-kb resolution intra-chromosomal matrices[58]. Hi-C matrix (M) was converted to correlation matrix C using corr (log2[M]). Then, the eigenvalues (λi) of matrix C was obtained by eigen-decomposition and normalized with 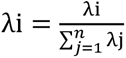. VNE was calculated as 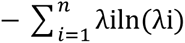

Compartments were called d by analyzing the first eigenvector of the KR normalized contact maps at 100 kb resolution. The compartments with higher gene density were assigned as type A, while the compartments with lower gene density were assigned as type B. Compartment strength was calculated using AB / AA + BB. Saddle plots were calculated as previously described[59]. Hi-C matrix bins were sorted according to the PC1 values. Sorted frequencies were aggregated into 50 groups and averaged to obtain a compartmentalization saddle plot. Number of ATAC-seq peaks overlapped with compartments were analyzed by BEDtools with at least 1bp shared[60].

TADs and TAD boundaries were identified at 50 kb resolution as described[61]. Shared TADs between different samples were defined as overlapping area larger than 75% for both samples. We calculated the standard deviations of the insulation score of each TAD boundary across three-cell stages and sorted boundaries by standard deviations. Then the top 1000 variable TAD boundaries were selected based on the ranking order. Clustering of boundaries was carried out using Pheatmap package in R.

Loops and interactions were detected with HiCCUPS in Juicer Tools at 10 kb resolution[62]. Enhancer or promoter involved loops were those with at least one anchor overlapped with enhancer regions (distal H3K27ac ChIP-seq peaks) or promoter regions (TSS ± 3 kb). Enhancer–promoter (E–P), promoter–promoter (P–P), and enhancer–enhancer (E–E) interactions were also identifed. Shared loops were defined as loops with both anchors not shift more than one bin. Aggregate peak analysis was processed with ‘apa’ in Juicer Tools, which generated aggregate heatmaps and average contact signals.

### TF motif analysis

Motifs were identified in H3K27ac ChIP-seq peaks located in loop anchors and the differential ATAC-seq peak regions using findMotifsGenome.pl in Homer[63]. The parameters were set as “-size given”. Only those motifs whose q-values smaller than 0.01 were treated as significantly enriched motifs.

### HPC7 analysis

ChIP-seq peaks of transcription factor, FPKM value of RNA-seq and loops identified from pcHi-C was downloaded from public data (GSE48086, GSE22178, E-MTAB-3954)[25, 27, 28]. Fraction of ChIP-seq peaks overlapping with loop anchors were compared with fractions of peaks overlapping with equal numbers of random chosen regions having same length with loop anchors. Genes were classified as PU.1 occupancy on both anchors if there is at least one loop connecting the gene promoter and having both anchors bound by PU.1. Genes with FPKM values ≥ 2 were considered as expressed.

## Authors’ contributions

WQZ and FFL conceived and designed the experiments. MH and XLL performed the experiments, MH, FFL, BXX, YBL and HKL analyzed the data. MH, FFL and ZYA wrote the paper. WQZ and FFL revised the manuscript. All authors read and approved the final manuscript.

## Competing interests

The authors have declared no competing interests.

## Acknowledgements

This work was supported by the Guangdong Natural Science Foundation (2023A1515030068, to Feifei Li), and the National Nature Science Foundation of China (grant no. 82270121, to Wenqing Zhang).

---

**Figure S1.**
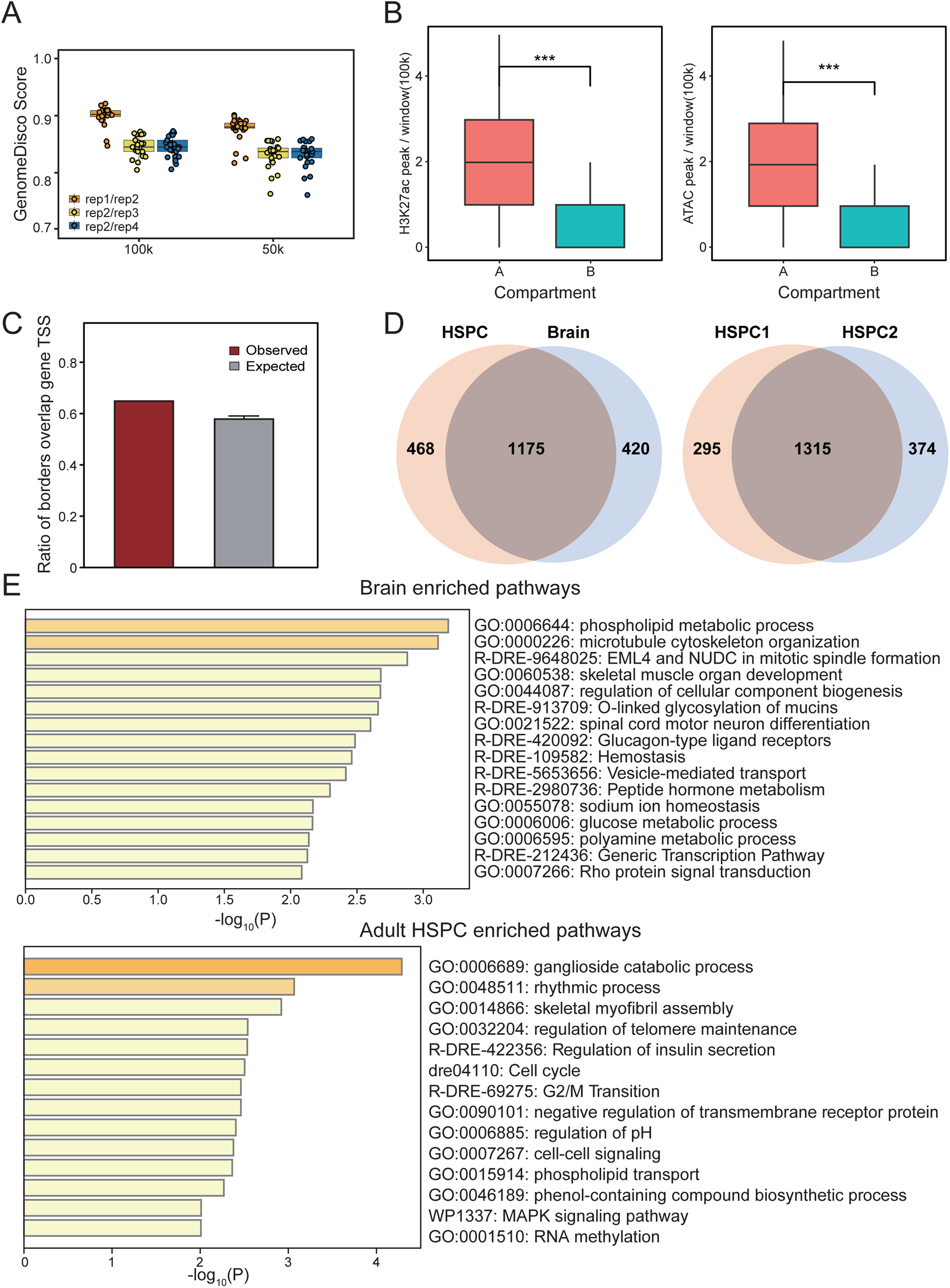
Chromatin conformation of zebrafish adult HSPC. (A) GenomeDisco scores showing the reproducibility of the Hi-C libraries. (B) Distribution of H3K27ac ChIP-seq and ATAC-seq peak density in the A/B compartments. (C) Boxplot showing proportion of TAD boundaries overlapped with gene transcriptional start site (TSS) and compared with random selected regions having same length as TAD boundaries. (D) Venn graphs showing the overlap of TADs between adult HSPC and brain as well as between biological replicates of HSPC. (E) The enriched biological processes of genes located in tissue-specific TAD boundaries.

**Figure S2.**
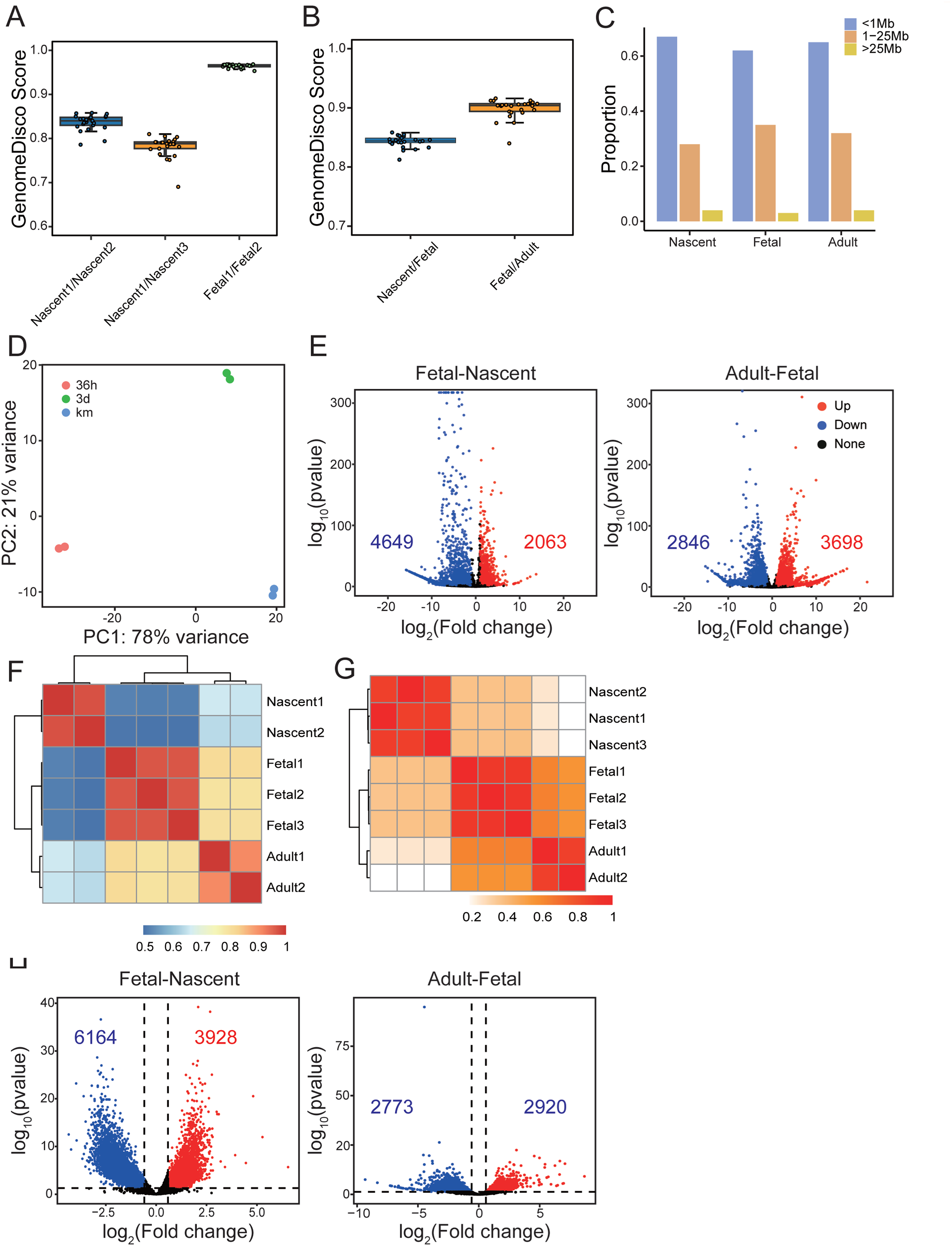
Global transcriptional and chromatin conformation changes during HSPC development. (A) GenomeDisco scores showing the reproducibility of the Hi-C libraries. (B) GenomeDisco scores of consecutive stages during HSPC development. (C) Proportion of interactions within different distances to all intra-chromosomal interactions. (D) Principal components analysis (PCA) of RNA-seq expression data for different samples. (E) Volcano plot of differentially expressed genes (Fold change ≥ 2, Padj < 0.05) for consecutive stages during HSPC development. (F) Clustering analysis of gene expression data for different samples. (G) Clustering analysis of ATAC-seq signal on accessible peak regions. (H) Volcano plot of differentially accessible regions based on ATAC-seq (Fold change ≥ 1.5, Padj < 0.05) for consecutive stages during HSPC development.

**Figure S3.**
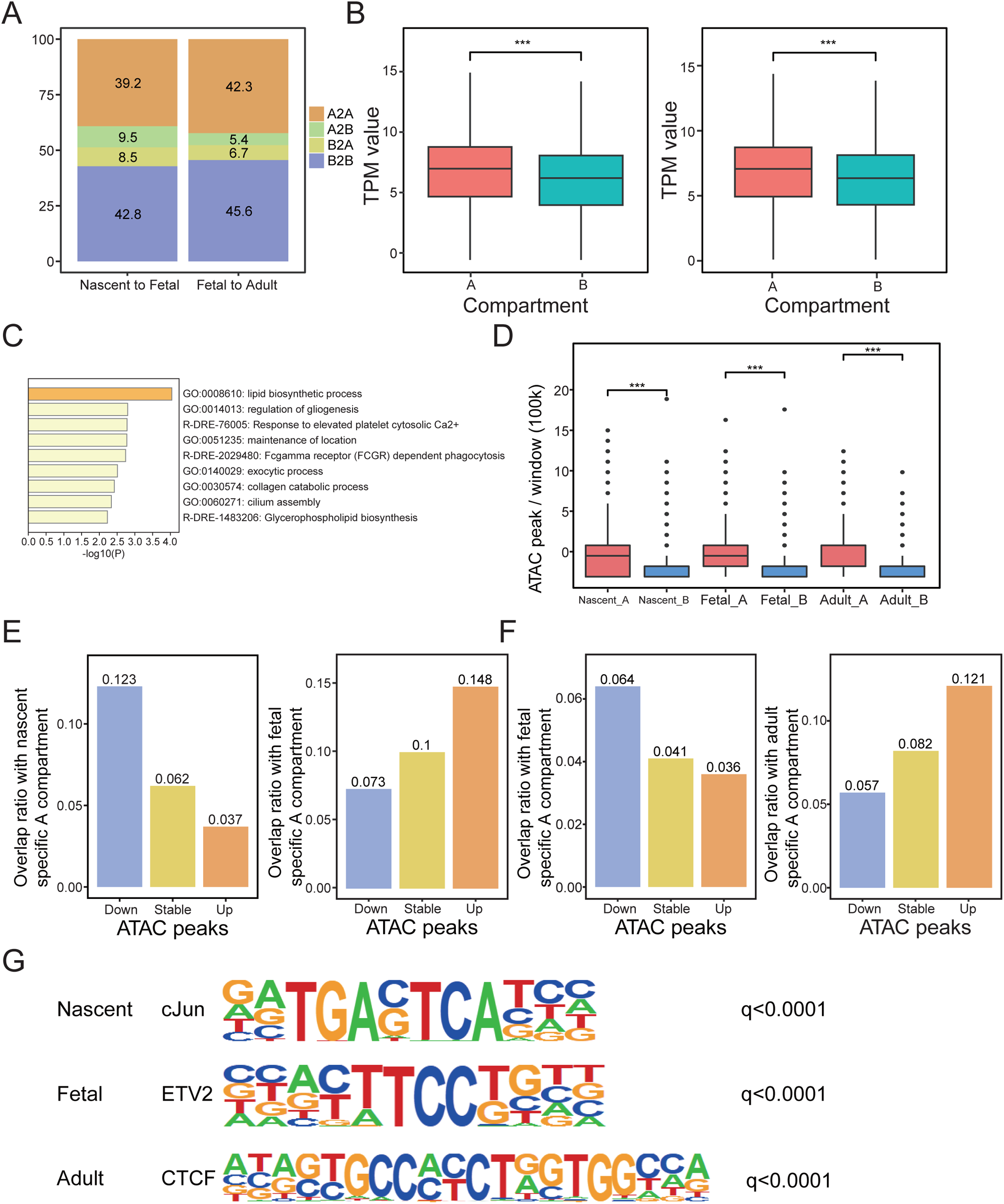
Changes of compartmentalization during zebrafish HSPC development. (A) Proportions of genome regions which switched compartments. (B) Boxplot showing the distribution of transcript per million (TPM) expression value of genes in the A/B compartment for nascent and fetal HSPC. (C) Enriched pathways of genes located in B2A switch region and upregulated from fetal to adult HSPC. (D) Distribution of ATAC-seq peak density in the A/B compartments for all developmental stages. (E) Overlap of differentially regulated ATAC-seq peaks from nascent to fetal HSPC with stage-specific compartment A regions. (F) The same as E, but from fetal to adult HSPC. (G) Most enriched TF and its motif within stage-specific accessible regions.

**Figure S4.**
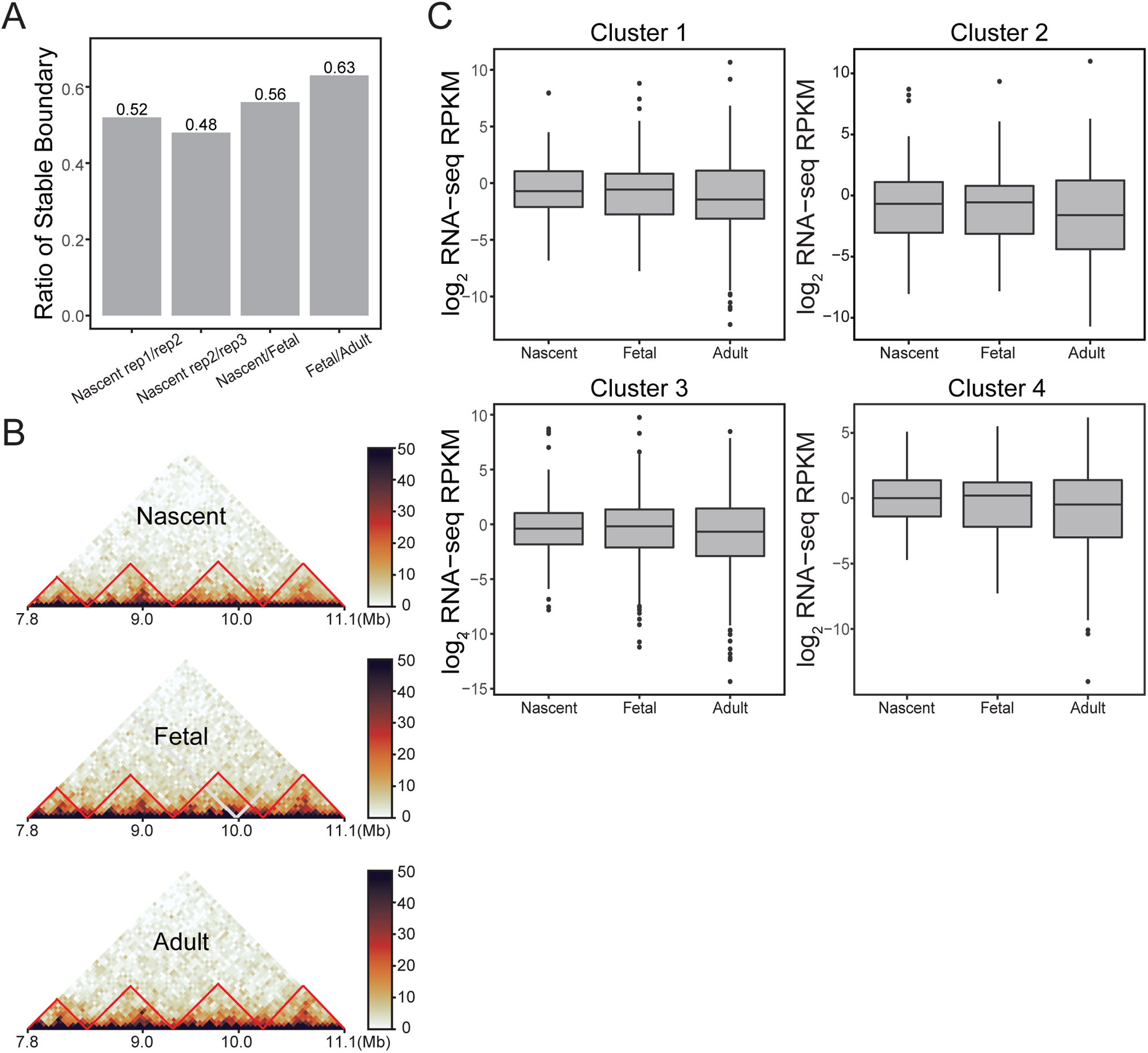
TADs kept relatively stable during zebrafish HSPC development. (A) Bar plot showing overlap of TAD boundaries between biological replicates and successive developmental stages. (B) Position of TADs in chr1: 7.8-11.1Mb for the three stages of HSPC. (C) Expression of genes located in different TAD boundary clusters.

**Figure S5.**
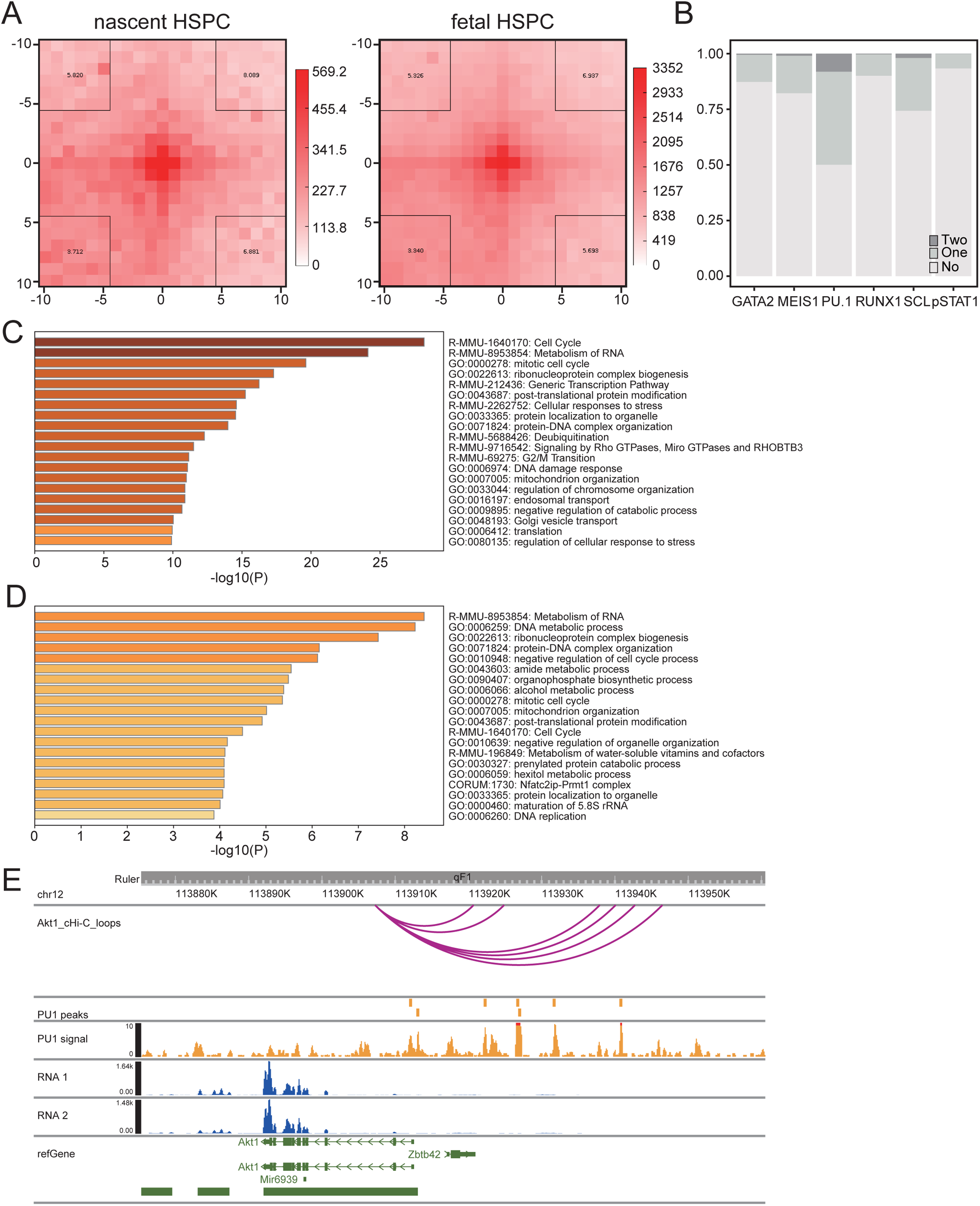
Candidate transcription factors mediating chromatin looping interactions in adult HSPC. (A) Aggregate loop plots showing contact frequencies of adult HSPC-specific loops in nascent and fetal HSPC cells. (B) Frequency of HPC7 loops occupied by each transcription factor. (C-D) Gene ontology analysis for genes having loop connecting gene promoter with only one anchor (C) or no anchor (D) occupied by PU.1. (E) Same as 5H, but near Akt1 gene.

