## Supplemental Table 1 for "Reprogramming of 3D genome structure underlying HSPC development in zebrafish"

| **Hi-C** | | | | | | | |
| --- | --- | --- | --- | --- | --- | --- | --- |
| **Samples** | **Total clean reads** | **Non-redundant valid pairs** | **reps valid pairs** | **Inter-chromosomal** | **Intra-chromosomal** | **Intra-chromosomal <20kb** | **Intra-chromosomal >20kb** |
| Nascent_rep1 | 392903427 | 8468150 | 26350389 | 4306214 | 4161936 | 935132 | 3226804 |
| Nascent_rep2 | 376885191 | 10849772 |  | 4619405 | 6230367 | 1481677 | 4748690 |
| Nascent_rep3 | 1129245164 | 7032467 |  | 4348858 | 2683609 | 892219 | 1791390 |
| Fetal_rep1 | 405830178 | 55235310 | 118555641 | 18525258 | 36710052 | 8181747 | 28528305 |
| Fetal_rep2 | 412265820 | 63320331 |  | 21721394 | 41598937 | 8824443 | 32774494 |
| Adult_rep1 | 383670036 | 45773503 | 172842225 | 12209492 | 33564011 | 9185550 | 24378461 |
| Adult_rep2 | 577041900 | 53946417 |  | 13988234 | 39958183 | 11322537 | 28635646 |
| Adult_rep3 | 248585257 | 35918086 |  | 12698822 | 23219264 | 6242842 | 16976422 |
| Adult_rep4 | 251625430 | 37204219 |  | 13272933 | 23931286 | 6182244 | 17749042 |

**Table S1**

| **Adult HSPC** | | | |
| --- | --- | --- | --- |
| **Type** | **samples** | **total reads** | **narrowPeaks** |
| ChIP-seq | H3K27ac-rep1 | 55196510 | 18802 |
|  | H3K27ac-rep2 | 35671326 |  |
|  | H3K27ac-input | 36963612 |  |
| ATAC-seq | ATAC-rep1 | 56866073 | 16052 |
|  | ATAC-rep2 | 18762859 |  |
| RNA-seq | RNA-rep1 | 36988724 |  |
|  | RNA-rep2 | 33170724 |  |
|  | RNA-rep3 | 38739195 |  |
